# Estrogen-dependent development and transcriptome regulation of the lateral septal kisspeptin system

**DOI:** 10.1101/2023.09.20.557932

**Authors:** Soma Szentkirályi-Tóth, Balázs Göcz, Szabolcs Takács, Miklós Sárvári, Imre Farkas, Katalin Skrapits, Éva Rumpler, Szilárd Póliska, Gábor Wittmann, Csaba Fekete, Erik Hrabovszky

## Abstract

While hypothalamic kisspeptin (KP) neurons play well-established roles in puberty and reproduction, kisspeptin neurons in the lateral septum (KP_LS_ neurons) and other extrahypothalamic sites have received less attention. We found that the onset of LS kisspeptin expression was linked to pubertal development and estrogen receptor signaling. Cell numbers were higher in females *vs.* males, continued to increase in adulthood and exogenous estrogen administered to adult mice was able to switch on the *Kiss1* gene promoter in new sets of septal neurons. Using RNA-Seq studies of laser-microdissected neurons from ovariectomized mice treated with 17β-estradiol (E2) *vs*. vehicle, we found that KP_LS_ neurons largely differ from hypothalamic KP neurons in their transcriptome profile which included 571 estrogen-dependent transcripts from which 80% were upregulated by a 4-day E2-treatment of ovariectomized mice. Notably, *Kiss1* expression in the LS was considerably lower than in hypothalamic KP neurons, being undetectable in ovariectomized mice and inducible by E2 supplementation. Finally, immunohistochemical detection of septal kisspeptin neurons and their fibers in the human brain suggested that the functions of this neuronal system are evolutionarily conserved. Ontogeny, sexual dimorphism and robust estrogenic regulation raise the intriguing possibility that the KP_LS_ system is a new central player in the estrogen-dependent control of reproductive and/or non-reproductive functions in mice, with a possible human relevance supported by the immunohistochemical observations on *post mortem* tissues.

## Introduction

Kisspeptin (KP) encoded by the *Kiss1* gene plays critical roles in the central regulation of puberty and fertility and inactivating mutations of *Kiss1* or the KP receptor gene *Kiss1r* cause hypogonadotropic hypogonadism in humans (1, 2) and mice (2-5). Conversely, activating mutations of these genes may account for clinical cases of precocious puberty (6, 7). Hypothalamic KP neurons promote fertility via stimulating luteinizing hormone (LH) secretion (8-11). This effect is mediated by gonadotropin-releasing hormone (GnRH) neurons which express *Kiss1r* (12, 13). Accordingly, deletion of *Kiss1r* from GnRH neurons causes infertility and its selective re-introduction to GnRH neurons in global *Kiss1r* knockout mice reinstates fertility (14).

Immunohistochemical and *in situ* hybridization studies localized the majority of KP synthesizing neurons to two main regions in the preoptic area/rostral hypothalamus (RP3V) and the hypothalamic arcuate nucleus (ARC), respectively (15). Both populations integrate and communicate to the GnRH system various external as well as internal cues, the latter including the positive and negative feedback effects of estrogens (16, 17). The KP cell group of the RP3V (KP_RP3V_ neurons) is formed by two subpopulations in the anteroventral periventricular nucleus (AVPV) and the rostral periventricular nucleus (18). These neurons are heavily implicated in the positive feedback effect of E2 triggering the proestrus afternoon surge of LH (16-18). They innervate GnRH neurons directly (19-21), express the activity marker c-Fos during the LH surge (22, 23) and respond to E2 with a series of transcriptional changes including an increased expression of *Kiss1* (24, 25).

The largest population of KP neurons is found in the arcuate nucleus (ARC) of the mediobasal hypothalamus which has long been known as a major sex steroid feedback site. Estrogen deficiency in postmenopausal women causes profound morpho-functional changes of KP_ARC_ neurons, characterized by neuronal hypertrophy (26, 27) and increased neurokinin B (NKB) (27, 28), KP (27, 29) and substance P (28, 30) biosynthesis. KP_ARC_ neurons (aka ‘KNDy’ neurons) also synthesize NKB and dynorphin in various mammals. They can stimulate GnRH release by a direct action of KP at GnRH nerve terminals (31), which plays an important role in negative sex steroid feedback (17, 32) and in the regulation of pulsatile GnRH/LH secretion (33).

While the role of hypothalamic kisspeptin neurons in the regulation of puberty and reproduction has been studied extensively in the past two decades, extrahypothalamic KP cell groups identified in the medial amygdala (MeA), the bed nucleus of the stria terminalis (BST) and the lateral septum (LS) have attracted relatively little attention (15, 34-39).

In the present study, we show that KP expression in lateral septal KP (KP_LS_) neurons is linked to pubertal development and estrogen receptor signaling. Cell numbers are sexually dimorphic and higher in females than in males. Exogenous estrogen and testosterone, but not dihydrotestosterone, turns on the *Kiss1* gene promoter in new sets of neurons. Using RNA-Seq studies of laser-microdissected KP_LS_ neurons from ovariectomized mice treated with 17β-estradiol (E2) *vs.* vehicle, we identify the unique transcriptome of KP_LS_ neurons which contains 571 estrogen-regulated transcripts. We show that septal *Kiss1* expression is undetectable in ovariectomized mice and induced by E2 replacement. Altogether, the ontogeny, sexual dimorphism and robust estrogenic regulation indicate that the KP_LS_ system is a new central player in the estrogen-dependent regulation of reproductive and/or non-reproductive functions in mice. Septal KP neurons and their fibers are also detectable in the human brain, raising the possibility of an evolutionarily conserved role.

## Results

### KP_LS_ neurons start to emerge during pubertal development and reach higher numbers in females

We first studied the postnatal ontogeny of KP_LS_ neurons in KP-Cre/ZsGreen mice. In this model, the *Kiss1* promoter driven expression of CRE recombinase switches on transcription of the fluorescent marker transgene in the ROSA 26 locus (36). Male and female mice from eight different age groups (N=4-7between weaning (PND 20-24) and PND 180-200 were perfused transcardially with 4% formaldehyde and their brains sectioned serially with a freezing microtome between plates 18-31 of the Paxinos mouse brain atlas (40). The number of KP_LS_ neurons was analyzed using fluorescent microscopy. We found that scattered KP neurons occurred first at PNW 5 in female and PNW 6 in male mice in the intermediate and dorsal subnuclei of the LS. Cell numbers increased gradually during and after pubertal development to reach a plateau of ∼ 90 in PND 180-200 males and of ∼180 in PND 140-160 females (**Figs. 1a, b**). In all age groups, KP_LS_ neurons were more numerous in females than in the age-matched males, with statistically significant sex differences (p<0.05; Mann-Whitney’s U-test) at PND 40-45 and later (**Figs. 1a, b**).

**Figure 1.**
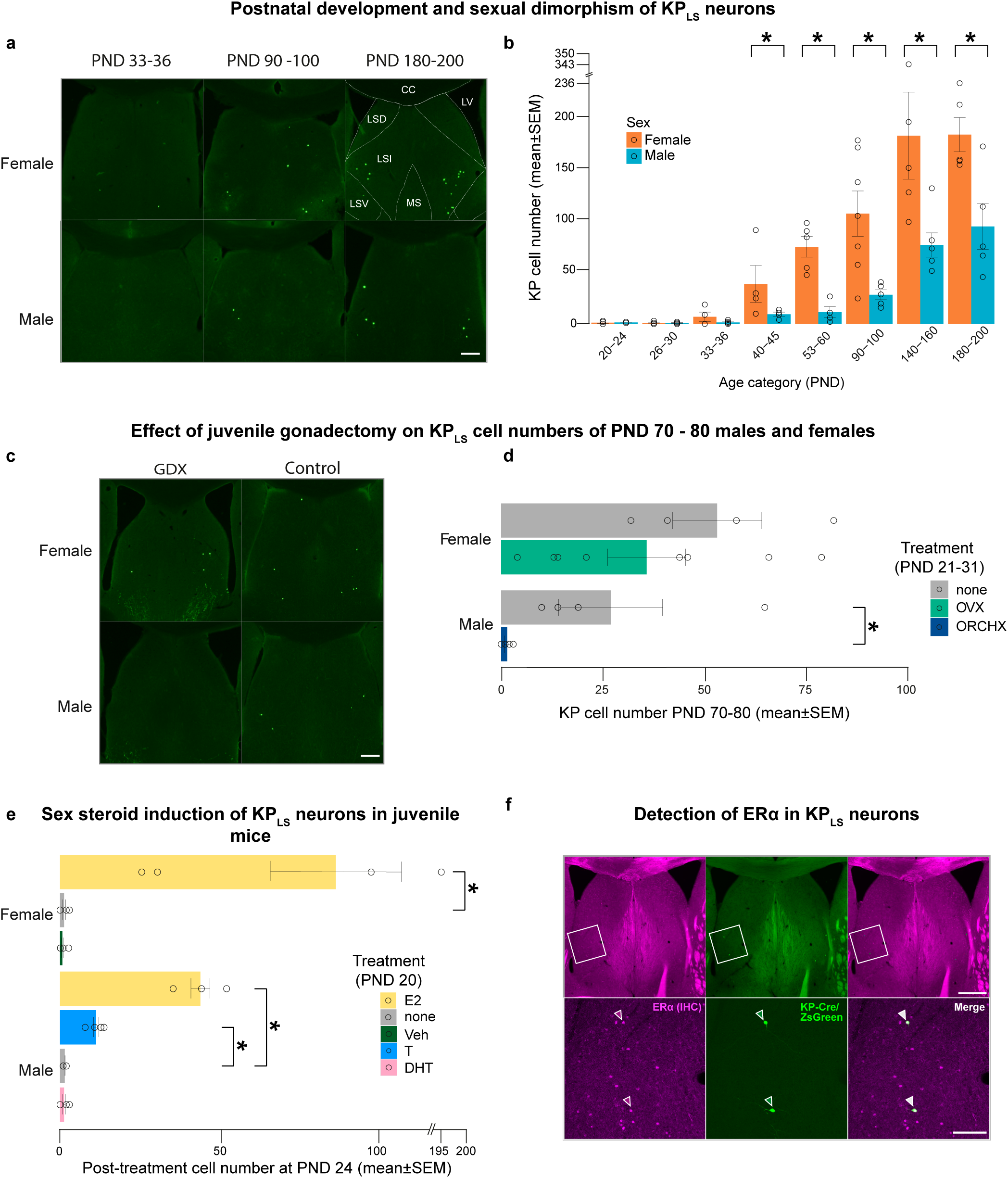
Developmental profile, sexual dimorphism and steroid induction of lateral septal kisspeptin neurons. **a:** Representative fluorescent photomicrographs illustrate various stages of KP_LS_ neuron development in male and female mice. **b:** Bar graphs reveal that KP-Cre/ZsGreen neurons begin to occur at PNW 5 in female and PNW6 in male mice and in all age groups, exhibit higher numbers in females (* p<0.05 by Mann-Whitney’s U-test). Cell numbers culminate at PND 140-160 in females and PND 180-200 in males. **c, d**: Adult PND 70-80 male mice orchidectomized (ORCHX) at PND 21-31 do not contain KP_LS_ neurons. In contrast, age-matched females ovariectomized (OVX) at PND 21-31 develop KP_LS_ neurons, although cell numbers tend to be somewhat lower than in age-matched intact controls. **e:** KP_LS_ neuron development can be advanced by a 4-day E2 treatment of juvenile female and male mice. Testosterone (T) also induces KP_LS_ neurons in male mice, albeit less efficiently. In this effect, T is likely converted to E2 to act on estrogen receptors, because the non-aromatizable androgen DHT is not able to induce KP_LS_ neurons. **f:** The immunohistochemical (IHC) localization of estrogen receptor-α (ERα; magenta) in the cell nuclei of KP_LS_ neurons (green) of adult female mice confirms that this classical estrogen receptor isoform plays a crucial role in the molecular regulation of KP_LS_ neurons. Scale bars: 250 µm in **a** and **c**; 400 µm in upper and 100 µm in lower panels of **f**. GDX: gonadectomy; LSD: lateral septum, dorsal subnucleus; LSI: lateral septum, intermediate subnucleus; LV: lateral ventricle; MS: medial septum.

### Developmental *Kiss1* expression in KP_LS_ neurons is linked to estrogen receptor signaling

This developmental profile together with the earlier evidence for the estrogenic upregulation of *Kiss1* mRNA expression in the LS (38) suggested that the prepubertal rise of gonadal sex steroids plays a role in KP_LS_ neuron development. Indeed, PND 70-80 male mice gonadectomized at PND 21-31 to prevent the pubertal rise of sex steroids did not contain KP_LS_ neurons (**Figs. 1c, d**). In contrast, KP_LS_ neurons were detectable in PND 70-80 females ovariectomized at PND 21-31, although cell numbers tended to be somewhat reduced (35.9 ± 9.6; mean ± SEM) in comparison with the age-matched female controls (53.3 ± 11; p=0.37; **Figs. 1c, d**). In this context, it is interesting to note that a much earlier ovariectomy of female mice at PND 15 was similarly unable to entirely prevent the sex steroid dependent development of KP_RP3V_ neurons, although cell numbers dropped by 70-90 % (41). To provide direct evidence for the role of sex steroids in the developmental induction of *Kiss1* expression, we implanted PND 20 mice (N=4) with a subcutaneous silastic capsule (l=10 mm; id=1.57 mm; od=3.18 mm) containing 100 μg/ml E2 in sunflower oil for a 4-day treatment. Control animals (N=4) received implants filled with vehicle. Exogenous E2 induced fluorescent signal in ∼80-90 septal neurons, mimicking the intact PND 90-100 female KP_LS_ cell numbers (**Fig. 1e**). Silastic capsules releasing E2 also induced KP_LS_ neurons in PND 20 males (N=3) by PND 24. The observation that the KP_LS_ cell numbers in E2-treated juvenile females were twice that of males, after the same treatment regime, raised the intriguing possibility that sexual dimorphism of KP_LS_ cell numbers might be imprinted earlier in life. Silastic implants (l=10 mm; id=1.47 mm; od=1.96 mm) filled with the male sex hormone T induced significantly lower cell numbers in male mice (N=4) than E2 (p=0.016 by Student’s t-test). As T can be partly converted to E2 by aromatase and thus, exert combined effects on estrogen and androgen receptors, the role of androgen receptor was addressed using another group of male mice (N=4) implanted at PND 20 with a silastic capsule (l=10 mm; id=1.47 mm; od=1.96 mm) releasing the non-aromatizable androgen dihydrotestosterone (DHT). DHT administration for 4 days did not induce fluorescent KP_LS_ neurons, indicating that the sex steroid induction of KP_LS_ neurons in males is mediated via ER, similar to females (**Fig. 1e**). Our studies suggest that estrogen signaling via its receptor(s) plays a critical role in KP_LS_ neuron development. Indeed, we have established that many ZsGreen-tagged KP_LS_ neurons express immunohistochemical signal for the classical estrogen receptor isoform, ERα (**Fig. 1f**).

### KP_LS_ neurons express a unique transcriptome profile and low levels of *Kiss1* mRNA

A laser capture microdissection/RNA-sequencing (‘LCM-Seq’) method developed in our laboratory (42) has enabled us to study and identify the estrogen-regulated transcripts of KP_ARC_ (43) and KP_RP3V_ (25) neurons. Here we characterized the transcriptome landscape of KP_LS_ neurons using the same animal models and protocols to allow the comparative analysis of the different KP cell types.

In brief, surgically ovariectomized adult (PND 90-110) KP-Cre/ZsGreen mice were implanted on post-ovariectomy day 9 with a single silastic capsule (l=10 mm; id=1.57 mm; od=3.18 mm) containing either E2 (100 μg/ml E2 in sunflower oil; OVX+E2 group; N=12) or sunflower oil vehicle (OVX+Veh group; N=9) for a four-day treatment. The brains were fixed with 0.5% formaldehyde followed by 20% sucrose using transcardiac perfusion. The LS was sectioned serially with a cryostat and ZsGreen-tagged KP neurons were isolated with LCM (25, 43). As the low number of KP_LS_ cells in individual mice may result in suboptimal detection sensitivity and quantification precision/accuracy of the LCM-Seq protocol (42), ∼360 fluorescently-tagged neurons were pooled proportionally from 3 brains to prepare each RNA sample and cDNA library, followed by Illumina sequencing (25, 43). Meta-analysis of the two hypothalamic KP neuron transcriptomes was carried out from open-access public repositories associated with the original publications (25, 43).

A marked difference between KP_LS_ neurons and the two hypothalamic KP cell types was observed in the level of *Kiss1* expression which was undetectable in KP_LS_ neurons of OVX+Veh mice and induced by E2 treatment in all four OVX+E2 samples to reach a mean CPM of 7.0. In contrast, *Kiss1* levels were considerably higher in both KP_ARC_ (CPM_OVX+Veh_: 570.5; CPM_OVX+E2_: 36.3) and KP_RP3V_ neurons (CPM_OVX+Veh_: 95.5; CPM_OVX+E2_: 777.5) (**Fig. 2b; Table S1**). Differential expression analysis with DeSeq2 identified thousands of other transcripts that were expressed differentially between KP_LS_ neurons and KP_ARC_ or KP_RP3V_ neurons either in the OVX+Veh or the OVX+E2 models (**Fig. 2c; Table S1**). For each KP cell type, the top 20 (with the highest mean CPMs, considering all OVX+Veh and OVX+E2 samples) transcription factors, transporters, ion channels and receptors (defined in the KEGG BRITE database) were selected and studied in all three cell types (**Table S2**). The top 20 lists combined from the three KP cell types included 31 ion channels, 28 transporters, 32 transcription factors, 29 neuropeptides and 31 receptors with little similarity in expression levels and regulatory patterns (**Fig. 2d; Table S2**).

**Figure 2.**
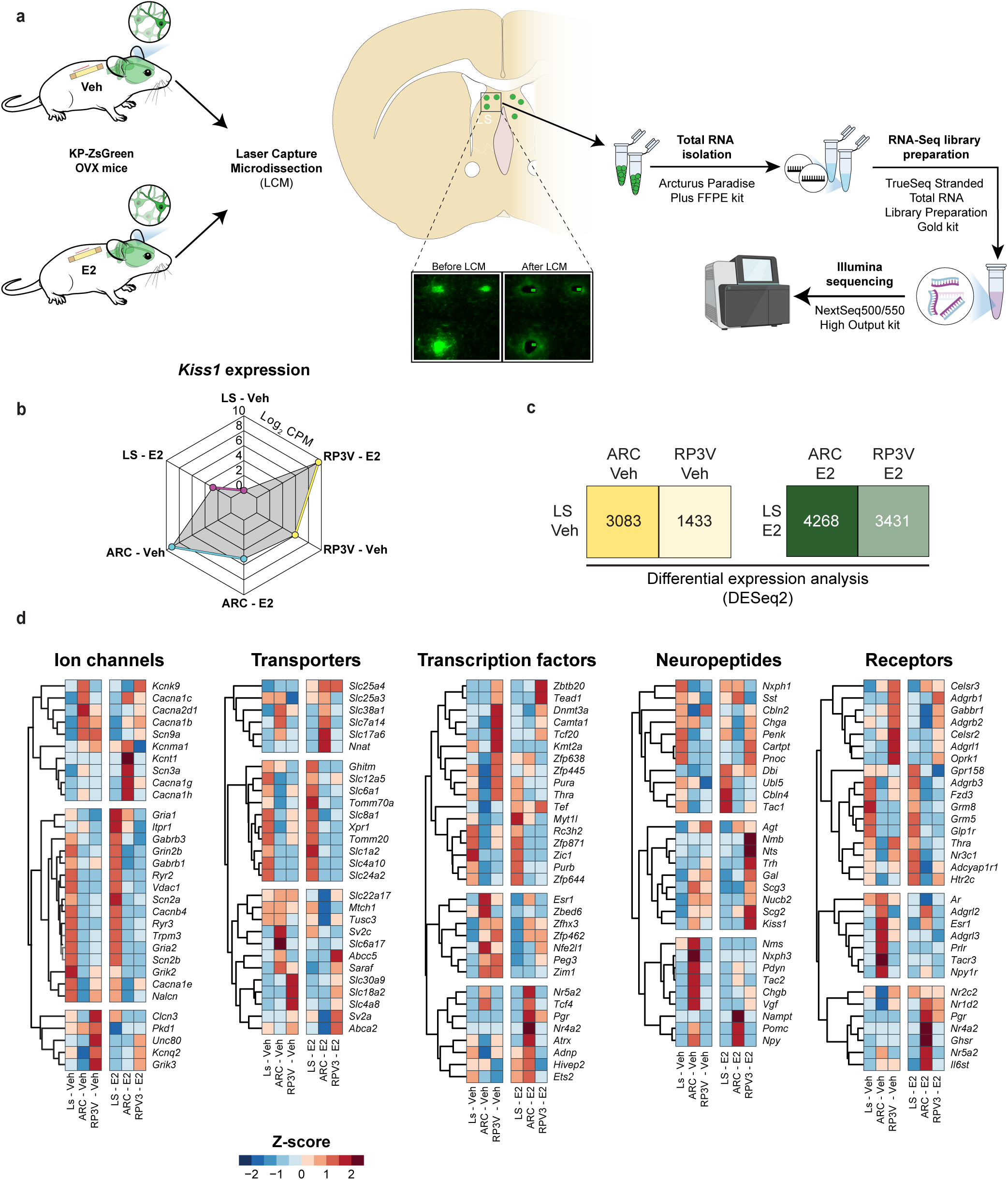
Transcriptome analysis with LCM-Seq reveals a unique gene expression profile of KP_LS_ neurons. **a:** Use of the OVX+Veh and OVX+E2 animal models and the LCM-Seq protocol to investigate the transcriptome profile of KP_LS_ neurons in the present and KP_ARC_ and KP_RP3V_ neurons in earlier gene expression studies allows comparison of the three distinct KP cell types. **b:** Spider chart illustrates that *Kiss1* expression is upregulated in KP_LS_ and KP_RP3V_ neurons and downregulated in KP_ARC_ neurons by E2 treatment. Notably, KP_LS_ neurons express considerably lower *Kiss1* levels than either KP_RP3V_ or KP_ARC_ neurons. **c:** Quantification table summarizes the number of genes expressed at significantly different levels (p.adj.<0.05 by DESeq2 analysis) between KP_LS_ neurons and the two hypothalamic KP cell types (see also **Table S1**). **d:** Heat maps compiled from the lists of the top 20 ion channels, transporters, transcription factors, neuropeptides and receptors of single KP cell types (**Table S2**) reveal large differences in expression levels (in CPM) and in patterns of estrogenic regulation.

### LCM-Seq unveils 571 estrogen-regulated transcripts in KP_LS_ neurons

Differential expression analysis of the OVX+Veh and the OVX+E2 transcriptomes with the DESeq2 R-package and RUVSeq normalization identified 571 transcripts which changed significantly (p.adj.<0.05) in estrogen-treated KP_LS_ neurons (**Fig. 3.; Table S3).** E2 upregulated ∼80% of the changing transcripts (**Figs. 3a, b)**. Overrepresentation (ORA) analysis identified significant changes in the Neuroactive ligand-receptor interaction, Glutamatergic synapse and Retrograde endocannabinoid signaling functional (KEGG) pathways (**Fig. 3c)**. Individual E2-regulated transcripts fell into diverse functional categories defined in KEGG BRITE and Neuropeptide (http://www.neuropeptides.nl/) databases, including ion channels (N=22), transporters (N=19), transcription factors (N=23), neuropeptides/granins (N=8), and receptors (N=15) (**Fig. 3d)**. For a full listing of E2-regulated KP_LS_ neuron transcript, see **Table S3**. Meta-analysis of the KP_ARC_ and KP_RP3V_ transcriptome databases with DESeq2 (using RuvSeq) identified 2458 and 547 transcripts, respectively, which were regulated significantly by the same E2 treatment (**Fig. 4; Table S3**). Comparison of the regulated transcripts in the three cell types confirmed the unique molecular properties and regulatory patterns of KP_LS_, KP_ARC_ and KP_RP3V_ neurons. Only 133 of the 571 E2-dependent KP_LS_ transcripts (∼23%) were regulated by E2 in the ARC and 58 (∼10%) in the RP3V (**Fig. 4**). In 46 instances, opposite regulation occurred in KP_LS_ neurons and either the KP_ARC_ or the KP_RP3V_ cell type. This phenomenon was reminiscent of several inverse regulatory changes we observed while comparing E2-induced changes in the KP_ARC_ and KP_RP3V_ cell types (25). E2-dependent transcripts of each KP cell types and overlaps between their E2-dependent transcripts are listed in **Table S3.**

**Figure 3.**
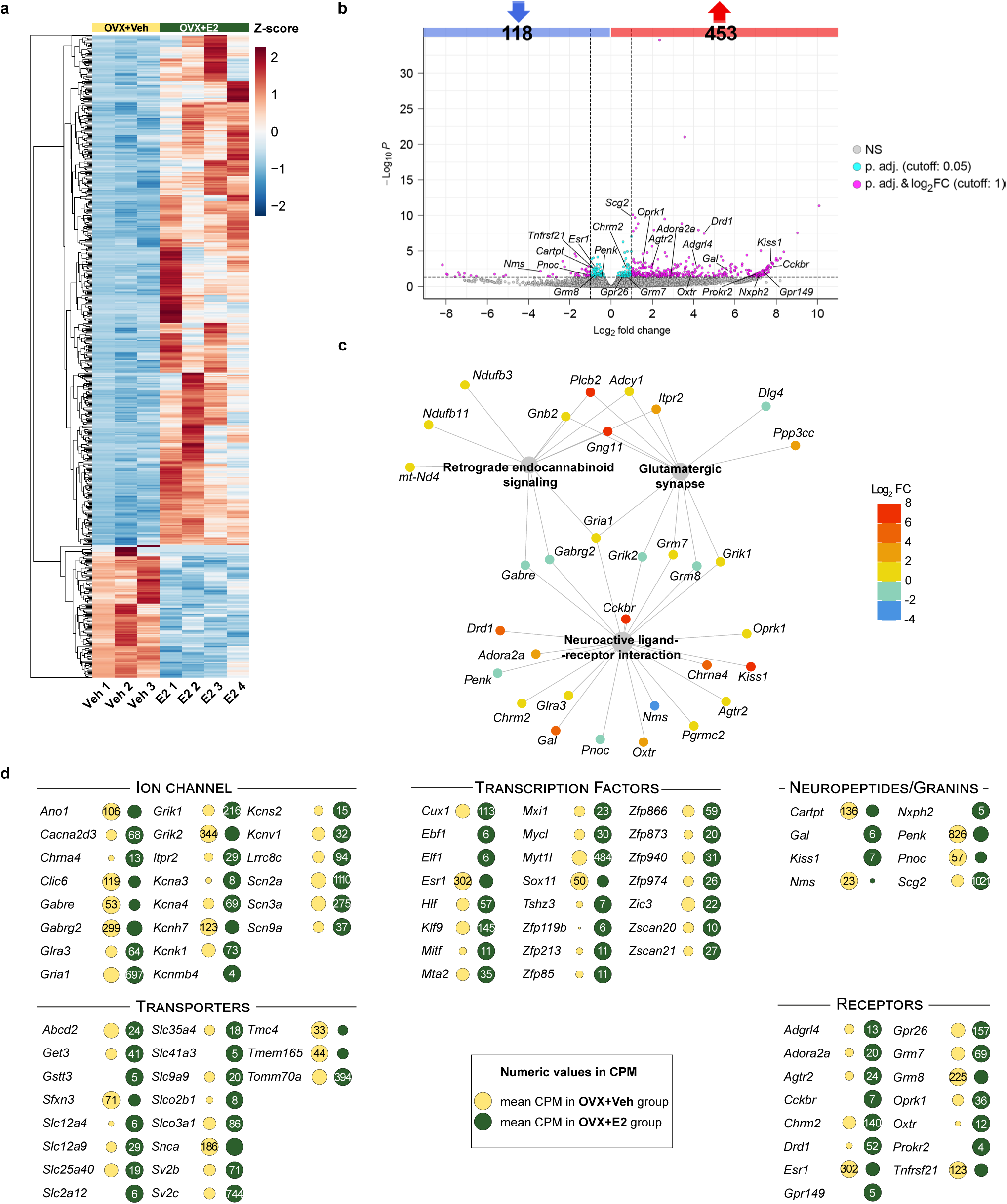
E2 treatment of OVX mice induces profound transcriptomic changes within KP_LS_ neurons. **a:** Heat map illustrates that the majority (∼80%) of the 571 transcripts altered by the 4-day E2 treatment are upregulated. (Dendrogram generated using UPGMA hierarchical clustering). **b:** Differentially expressed genes are shown in a volcano plot. Neuropeptides/granins and receptors are tagged with a gene symbol. **c:** Overrepresentation (ORA) analysis identifies three significantly altered functional (KEGG) pathways. E2-regulated transcripts of the Neuroactive ligand-receptor interaction, Glutamatergic synapse and Retrograde endocannabinoid signaling categories are illustrated graphically. **d:** Individual E2-regulated transcripts fall into diverse functional categories defined in KEGG BRITE and Neuropeptide databases. Many of them encode ion channels (N=22), transporters (N=19), transcription factors (N=23), neuropeptides/granins (N=8) and receptors (N=15). Numbers in dots refer to relative transcript abundances in CPM units. Dot areas change in proportion to transcriptional responses, with the larger dot representing 100%. See **Table S3** for full listing of E2-regulated KP_LS_ neuron transcripts.

**Figure 4.**
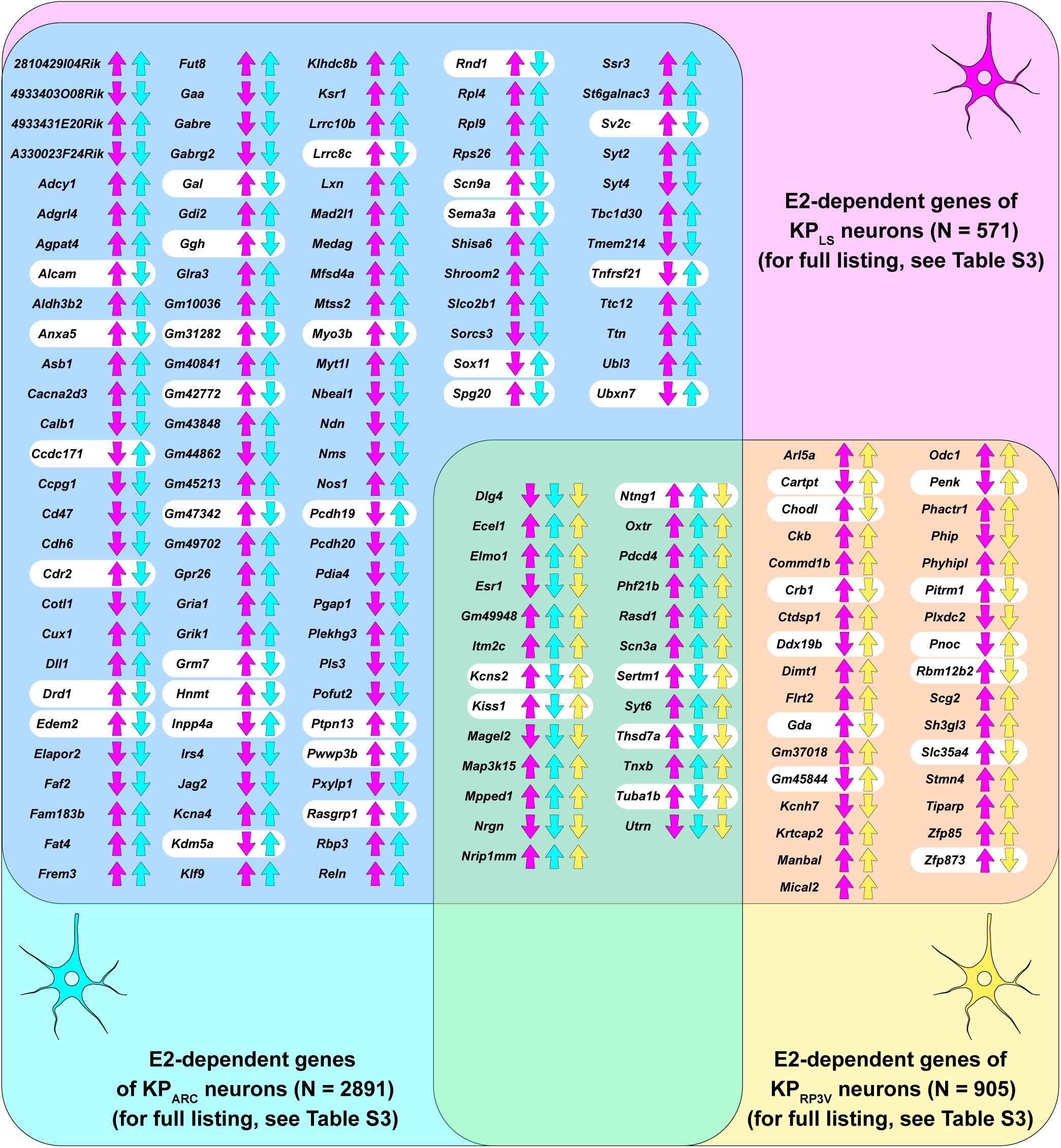
Distinct E2-dependent gene sets of LS, ARC and RP3V KP neurons indicate unique functional properties of the three distinct KP cell types. Venn diagram illustrates 166 out of the total 571 KP_LS_ neuron transcripts that are also regulated by E2 in KP_ARC_ and/or KP_RP3V_ neurons. For full listing of the regulated transcripts in each cell type, see **Table S3**. White labels highlight genes with inverse regulatory patterns.

### Estrogenic upregulation of the *Chrm2* transcript enhances muscarinic inhibition of KP_LS_ neurons

Muscarinic and nicotinic receptor transcripts in the KP_LS_ neuron transcriptome indicated that acetylcholine plays a role in the afferent control in KP_LS_ neurons. Indeed, fibers immunoreactive to the cholinergic marker vesicular acetylcholine transporter (VAChT) surrounded KP_LS_ neurons and established appositions to KP cell bodies and dendrites (**Fig. 5a**). From the muscarinic and nicotinic receptor forms detected in KP_LS_ neurons with RNA-Seq, *Chrm2* showed the highest expression level and a strong positive regulation by E2 (**Fig. 5b**), allowing us to predict an enhanced muscarinic inhibition of KP_LS_ neurons in OVX+E2 *vs.* OVX+Veh mice. To provide functional support to this hypothesis, the effect of muscarine on the spontaneous firing and resting membrane potential (Vrest) of KP_LS_ neurons were studied with whole-cell patch clamp electrophysiology (**Fig. 5c**). KP_LS_ neurons were spontaneously active, with phasic-irregular (∼ 90 %) or tonic (≈ 10 %) firing, a rate of 3.1 ± 0.55 Hz in OVX+Veh (n=15) and 2.9 ± 0.58 Hz (n=12) in OVX+E2 mice (not different statistically; Student’s t-test, p=0.8035) and a resting potential of -48 ± 1.3 mV in OVX+Veh and -48 ± 1.5 mV in OVX+E2 mice (not different; Student’s t-test, p=0.9008). A single bolus of muscarine (100 μM) transiently hyperpolarized and abolished the firing of KP_LS_ neurons which lasted longer in the OVX+E2 model (**Figs. 5d, e**). Longer inhibition was reflected in a larger decrease in the firing rate calculated for the 7-min period after muscarine treatment which dropped to 26.7 ± 6.58% of the control rate in OVX + E2 mice and to 54.8 ± 7.33% in OVX+Veh mice (**Table S4**). Addition of the M2 muscarinic acetylcholine receptor antagonist methoctramine (Methoct; 2 μM) to the aCSF entirely prevented the muscarine-induced hyperpolarization and silencing of KP_LS_ neurons, providing evidence for the major contribution of this receptor in muscarinic inhibition (**Fig. 5f**). Muscarine-induced hyperpolarization of KP_LS_ neurons in the OVX+E2 model persisted when action potentials were inhibited with 660 nM TTX (ΔVrest = -10.6 ± 2.00 mV), indicating that muscarine acted directly (**Fig. 5g**). OVX + E2 and OVX+Veh mice differed significantly (p<0.05; see **Table S4** for detailed statistics) in several electrophysiological measures which, together, provided support to the idea that the estrogenic induction of *Chrm2* expression causes heavier muscarinic inhibition in the previous model. These included the percentage of firing rate calculated for the 7 min post-treatment period (**Fig. 5h**; 54.8 ± 7.33% of the control rate in OVX+Veh and 26.7 ± 6.58% in OVX+E2 mice), the length of the silent period (**Fig. 5i**; 157 ± 28.4 s in OVX+Veh and 290 ± 30.3 s in OVX+E2 mice), and hyperpolarization amplitude **(Fig. 5j**; ΔVrest; -8.0 ± 0.98 mV in OVX+Veh and - 12.2 ± 1.14 mV in OVX+E2 mice).

**Figure 5.**
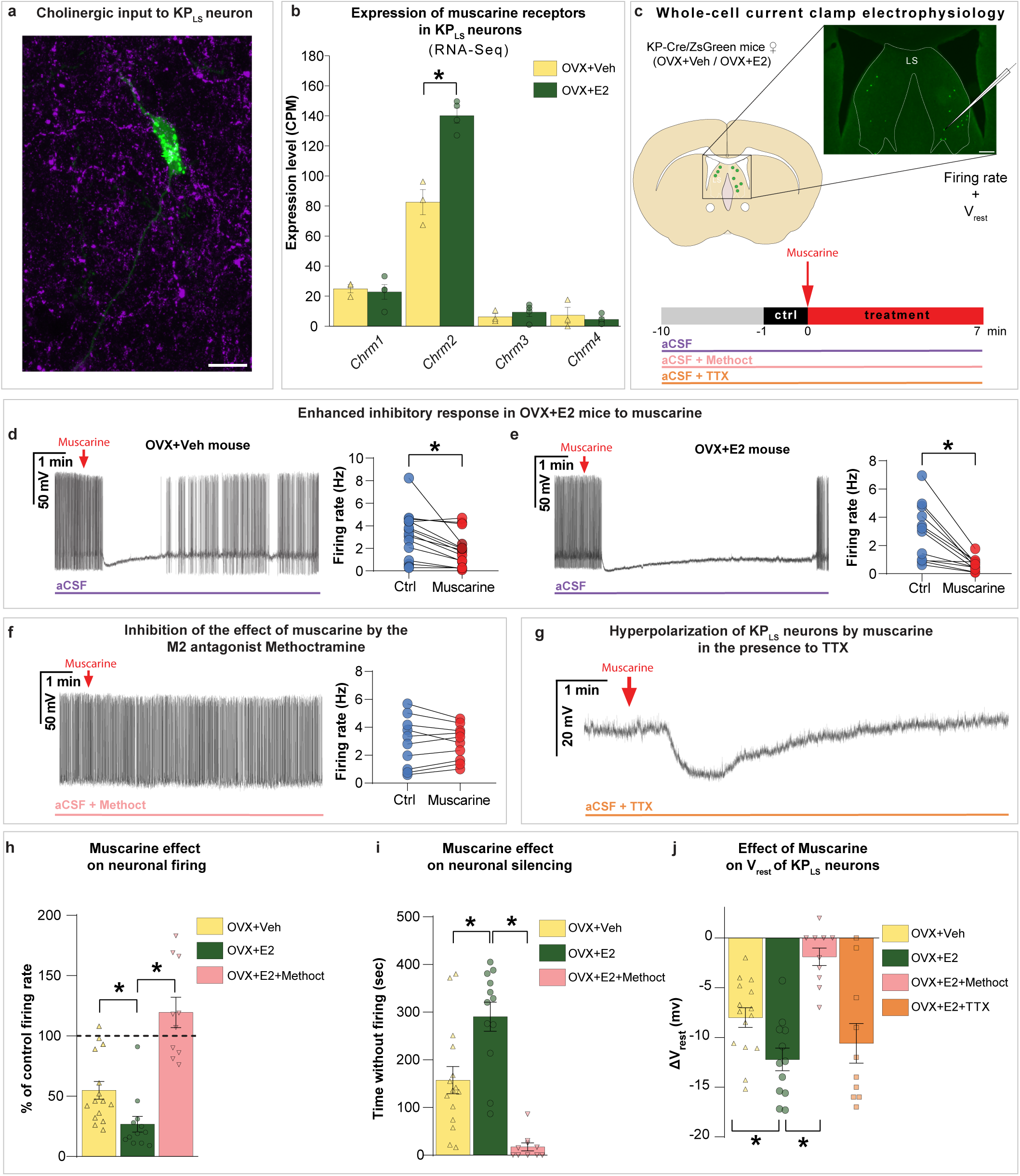
Estrogen-induced upregulation of the M2 acetylcholine receptor enhances muscarinic inhibition of KP_LS_ neurons. **a:** Confocal microscopic analysis reveals that KP_LS_ neurons (green ZsGreen signal) are embedded in a dense network of cholinergic fibers immunoreactive to the vesicular acetylcholine transporter (magenta). **b:** KP_LS_ neurons express several muscarinic acetylcholine receptors. High levels and estrogenic upregulation of *Chrm2* encoding for the inhibitory M2 muscarinic acetylcholine receptor suggest that the strength of cholinergic inhibition on KP_LS_ neurons is estrogen-dependent. **c:** Use of whole-cell patch-clamp electrophysiology allows the comparative analysis of muscarinic responses from OVX+Veh *vs.* OVX+E2 mice. **d, e**: Representative traces obtained from OVX+Veh and OVX+E2 mice illustrate that a single bolus of muscarine (100 μM) hyperpolarizes KP_LS_ neurons and transiently abolishes their firing. Note that the silent period lasts longer in the OVX+E2 model (**e**). The decreased firing rates shown in graphs were calculated for the 7-min period after muscarine administration. **f**: Presence of the M2 muscarinic acetylcholine receptor antagonist methoctramine (Methoct; 2 μM) in the aCSF medium prevents the muscarine-induced hyperpolarization and silencing of KP_LS_ neurons. **g**: Muscarine-induced hyperpolarization of KP_LS_ neurons persists in the presence of tetrodotoxin (TTX, 660 nM), suggesting that muscarine acts directly on these cells. **h, i**: Bar graphs reveal that the percentage decrease in the firing rates (**h**) and the length of the silent period (**i**) differ significantly between KP_LS_ neurons of OVX+E2 mice and OVX+Veh mice and between KP_LS_ neurons of OVX+E2 mice with and without Methoct pretreatment. **j**: Bar graph reveals that the amplitude of muscarine-induced hyperpolarization is significantly larger in OVX+E2 *vs.* OVX+Veh mice. Arrows indicate the time when muscarine is added to the aCSF. **\*=**p<0.05. For source data, see **Table S4.** Scale bars: 10 µm in **a** and 200 µm in **c**.

### Fluorescent *in situ* hybridization results confirm that KP_LS_ neurons use other peptides and gamma aminobutyric acid for neuronal communication with efferent targets

RNA-Seq studies revealed the expression of several neuropeptides and GABAergic markers in KP_LS_ neurons. We carried out fluorescent *in situ* hybridization (FISH) studies with digoxigenin-labeled cRNA probes and confirmed that KP_LS_ neurons synthesize the neuropeptide CART (encoded by *Cartpt*; **Fig. 6a**) and the GABAergic marker enzyme GAD1 (**Fig. 6b**).

**Figure 6.**
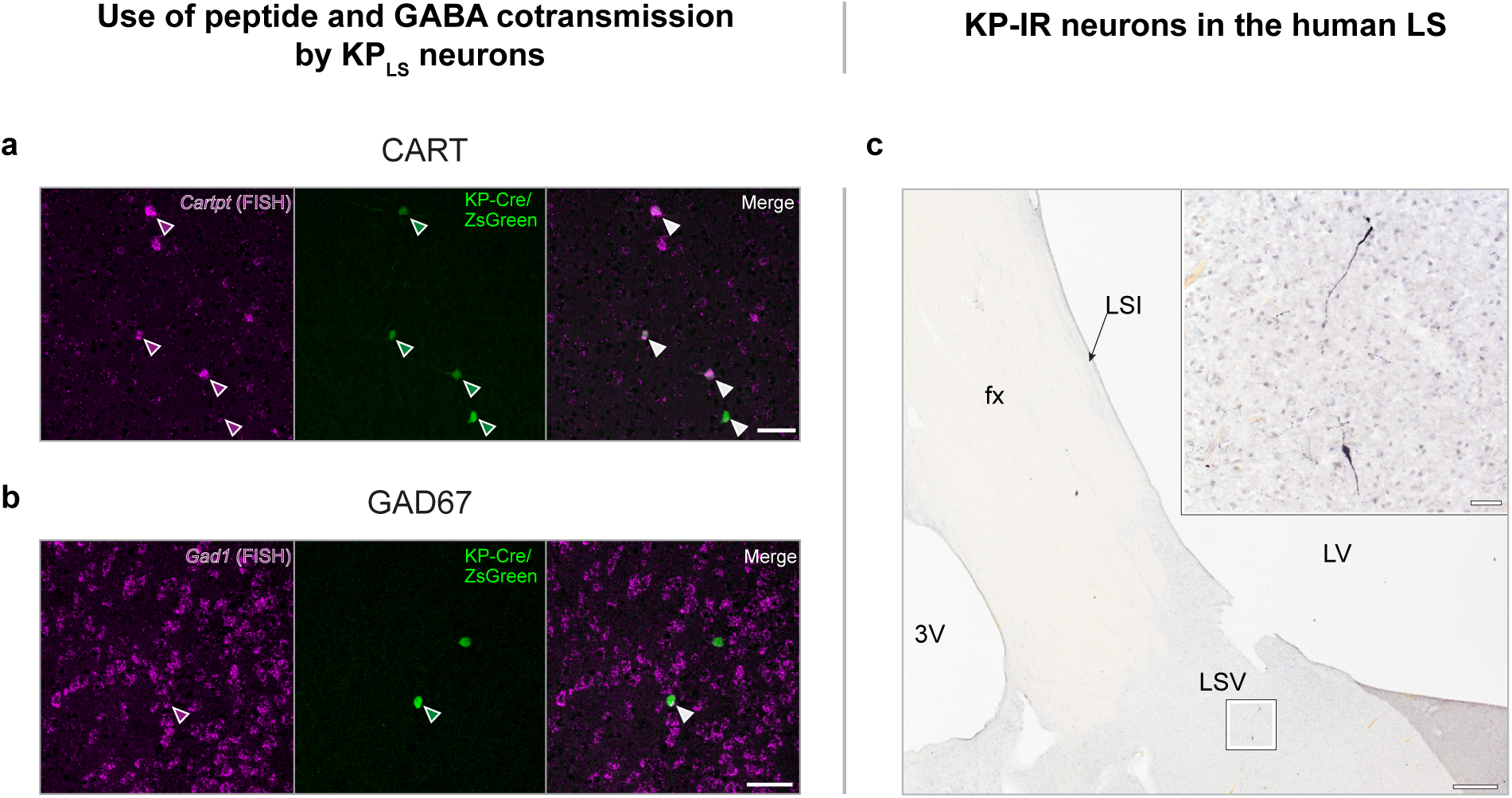
Neuroanatomical approaches reveal peptide and amino acid co-transmitters in murine KP_LS_ neurons and provide evidence for the presence of KP_LS_ neurons in the *postmortem* human brain. **a:** Confirming RNA-Seq results, fluorescent *in situ* hybridization (FISH) detection of *Cartpt* mRNA with a digoxigenin-labeled cRNA probe provides evidence that large percentages of KP_LS_ neurons use cocaine and amphetamine regulated transcript (CART; magenta) as a peptide co-transmitter. **b:** Co-expression of *Gad1* mRNA serves as a marker of GABAergic neurotransmission. Arrowheads in **a** and **b** point to KP_LS_ neurons (ZsGreen) that express the FISH signal. **c:** Immunohistochemical detection of KP (black nickel-diaminobenzidine chromogen) in the *postmortem* brain of a 53-year-old male subject provides evidence that KP neurons are also present in the human LS. Scale bars: 50 µm in **a**, **b** and in the high-power inset in **c** and 500 µm in the low magnification panel **c.** fx: fornix; LSI: lateral septum, intermediate subnucleus; LSV: lateral septum, ventral subnucleus; LV: lateral ventricle; 3V: third ventricle.

### Detection of kisspeptin neurons in the human septum suggests evolutionarily conserved roles of KP_LS_ neurons

In previous studies from our laboratory we provided evidence that the human hypothalamus contains the same two hypothalamic KP neuron populations that are observed in a variety of species. To determine if KP_LS_ neurons are present in the human septum, we carried out immunohistochemical studies on *postmortem* human septum samples (N=3) immersion-fixed with formaldehyde. Immunohistochemical detection of the human KP54 peptide revealed a few scattered KP immunoreactive neurons in the lateral septum of the human brain within a moderately dense KP fiber plexus (**Fig. 6c**). This observation suggests the intriguing possibility that the function of KP_LS_ neurons is evolutionarily conserved and likely plays a regulatory role in human septal functions.

## Discussion

In this study we report the estrogen-dependent development, sexual dimorphism and transcriptome profile of the KP_LS_ neuronal system in mice and also reveal homologous KP neurons in the LS of the *post mortem* human brain. These data indicate that KP_LS_ neurons exert estrogen-dependent and sex-specific control of currently unknown reproductive and/or non-reproductive functions in mice, with possible human relevances.

### Role of estrogens in the regulation of KP_LS_ neurons

Detection of *Esr1* mRNA expression and ERα immunoreactivity in KP_LS_ neurons in this study, and the robust E2-dependent regulation of the KP_LS_ neuron development and transcriptome, are in accordance with the results of earlier *in situ* hybridization studies which showed that E2 positively regulates *Kiss1* mRNA expression in this brain area (35, 38).

### KP neurons of the lateral septum possess a unique transcriptome profile

Earlier studies using transgenic mice expressing tdTomato reporter identified KP neurons outside the hypothalamus including the LS, the MeA, the periaquaductal grey and the mammillary nuclei (36). While much attention has been paid to the reproductive significance of the two hypothalamic KP neuron populations, functions of the extra-hypothalamic KP neurons are still relatively unexplored. Here we have used an LCM-Seq method developed in our laboratory (42) to study the transcriptome profile of KP_LS_ neurons and its estrogen-dependent regulation. This method allowed us to identify more than 15,000 transcripts which could then be compared to the transcripts of KP_ARC_ and KP_RP3V_ neurons we studied earlier using the same methodologies and animal models (25, 43).

The LS expresses high levels of glutamic acid decarboxylase *Gad1*, *Gad2* and AMPA glutamate receptors (44). Among calcium-binding protein markers, LS neurons express calbindin (45) or calretinin (46), but not parvalbumin. In accord with this general expression profile, we found that KP_LS_ neurons also show robust expression of *Gad1*, *Gad2*, *Calb1* and *Gria1*, and only low *Calb2* (calretinin) and *Pvalb* expression. KP_LS_ neurons exhibit a unique transcription factor profile, with several homeobox (*Meis2*, *Otx2*, *Lhx6*, *Lhx8*) and zinc finger (*Zic1-Zic5*) proteins, and some major transcription factors characteristic also of other KP cell types (e.g. *Myt1l*, *Esr1*).

### Comparative analysis of different KP cell types

Using differential expression analysis of RNA-Seq libraries from OVX+Veh and OVX+E2 mice, we have identified hundreds of E2-regulated transcripts in KP_LS_ neurons, including *Kiss1*. Upregulation of *Kiss1* by E2 was in agreement with the results of previous *in situ* hybridization studies by Stephens and colleagues (38). We found that E2 predominantly activated the expression of genes in KP_LS_ neurons, with a large number of upregulated transcripts related to cholinergic, dopaminergic and glutamatergic neurotransmission. Using whole-cell current-clamp electrophysiology, we provided functional evidence that the high expression of the *Chrm2* transcript in the OVX+E2 model enhances muscarinic inhibition of KP_LS_ neurons.

We have previously characterized the E2 regulated transcriptomes of KP_ARC_ (43) and KP_RP3V_ (25) neurons. These data together with the recently published elegant reports from the Kauffman and Stephens laboratories on the estrogenic regulation of the active KP_RP3V_ (24) and KP_MeA_ (39) neuron translatome allowed us to study the similarities and differences between various hypothalamic and extrahypothalamic KP cell types via the re-analysis of the deposited sequencing files using DESeq2 with RUVseq normalization. In general, each KP cell type showed very unique gene expression patterns, with conspicuous differences also in estrogenic regulation. We identified 2891, 905, 571 and 101 E2-regulated transcripts in KP_ARC_, KP_RP3V_, KP_LS_ and KP_MeA_ neurons, respectively, revealing that transcriptional responses to E2 are strongest in KP_ARC_ neurons and weaker in extrahypothalamic regions. An important difference in estrogen responses was the upregulation of *Pgr* in hypothalamic, but not in extra-hypothalamic KP neurons. Further re-analysis of the three datasets generated in our laboratory using the same methodologies, we determined transcriptional changes that overlapped and others that showed opposite patterns in KP_ARC_, KP_RP3V_ and KP_LS_ neurons. Based on these results we conclude that each KP neuron population exerts a unique transcriptional response to E2 via ERα with a great number of robust changes in different elements of the neuroactive ligand-receptor pathway.

### Sexual dimorphism

We have identified significantly higher KP_LS_ cell numbers in females *vs*. males both during development and in adults. We have also observed a similar sex difference when KP neurons were induced by the same steroid treatment of juvenile male and female mice, raising the possibility that sexual dimorphism is due, at least partly, to the organizational effect of an even earlier sex steroid exposure. Our data about the presence of KP_LS_ neurons in adult female, but not male, mice gonadectomized in the juvenile period strongly support this hypothesis.

Female dominance also characterizes KP neurons in the rodent RP3V (41, 47), in accordance with the phenomenon that only females exhibit positive estrogen feedback in rats and mice. Interestingly, *in situ* hybridization studies of *Kiss1* mRNA expression in the MeA revealed an opposite pattern of dimorphism, with higher cell numbers in intact male than in diestrus female mice (34). Similar *Kiss1* cell numbers in gonadectomized female and male rats treated with the same dose of E2, indicate that the sex difference in the MeA is due mostly to the dissimilar sex steroid milieu.

In conclusion, in this study we report the sex steroid-dependent ontogeny of KP_LS_ neurons in mice. We provide evidence that this system is sexually dimorphic, with a female dominance. We characterize the unique transcriptome profile of KP_LS_ neurons and describe its E2-dependent regulation and differences from other KP neuron populations. Finally, we show that a homologous neuron population is also present in the human brain. Further studies will be required to identify the efferent targets and roles of the KP_LS_ neuronal system in the regulation of reproductive and/or non-reproductive functions.

## Materials and Methods

### Animals

Animal experiments were carried out in accordance with the Institutional Ethical Codex, Hungarian Act of Animal Care and Experimentation (1998, XXVIII, section 243/1998) and the European Union guidelines (directive 2010/63/EU) and approved by the Animal Care and Use Committee of the Institute of Experimental Medicine. All efforts were made to minimize potential pain or suffering and to reduce the number of animals used. The mice (N=174) were housed under standard conditions (lights on between 0600 and 1800 h, temperature 22±1 °C, chow and water ad libitum) in the animal facility of the Institute of Experimental Medicine. KP-Cre/ZsGreen and KP-Cre/tdTomato mice were generated by crossing heterozygous Kiss-Cre (36) males with females of the Ai6 RCL-ZsGreen (B6.Cg**-***Gt(ROSA)26Sor^tm6(CAG-ZsGreen1)Hze^*/J) line (Jackson Laboratory, IMSR_JAX:007906) or the Ai14 (B6.Cg-*Gt(ROSA)26Sor^tm14(CAG-tdTomato)Hze^*/J) line (Jackson Laboratory, IMSR_JAX:007914), respectively.

### Quantitative analysis of KP_LS_ cell numbers with fluorescent microscopy

KP-Cre/ZsGreen mice (N=82) were anesthetized with a cocktail of ketamine (25 mg/kg), xylavet (5 mg/kg), and pipolphen (2.5 mg/kg) in saline and perfused transcardially with 80 ml of ice-cold 4% formaldehyde (prepared freshly from paraformaldehyde) in 0.1 M phosphate buffered saline (PBS; 0.1 M; pH 7.4). The brains were postfixed for 2 hours, infiltrated with a 20 % sucrose solution in PBS, snap-frozen in powdered dry ice and stored at -80 °C until sectioned at 40 µm in the coronal plane with a Leica SM 2000R freezing microtome (Leica Microsystems). The sections were stored at -20 °C in 24-well tissue culture plates filled with antifreeze solution (30% ethylene glycol, 25% glycerol, 0.05 M phosphate buffer (pH 7.4). To determine cell counts, all sections of the LS between plates 18-31 of the Paxinos mouse brain atlas (40) were analyzed with a Zeiss AxioImager M1 microscope (Carl Zeiss, Göttingen, Germany) using a 10× objective lens and a filter set for fluorescein isothiocyanate. Representative fluorescent images of the LS were captured with an AxioCam MRc5 digital camera (Carl Zeiss) using the AxioVision Se64 Rel.4.9.1 software. Editing was performed with Adobe Photoshop CS5 and the final composite images were created with Adobe Illustrator (Adobe System Inc., California, USA).

### Surgical treatments to generate mouse models with low and high physiological 17**β**-estradiol levels

The OVX+Veh and the OVX+E2 adult animal models were generated as described in previous studies (25, 43). In brief, the mice were surgically ovariectomized in ketamine (25 mg/kg)/xylazine (5 mg/kg)/promethazine (2.5 mg/kg) anesthesia. To prevent infections, Augmentin (GlaxoSmithKline plc, Brentford, UK; 30 µg/10 g bw in saline) was injected into the peritoneal cavity during the surgery and Baneocin ointment (Sandoz Gmbh, Kundl, Austria) was applied to the wound before the skin was clipped. On postovariectomy day 9, the animals were re-anesthetized and implanted subcutaneously with a single silastic capsule (Sanitech, Havant, UK; l=10 mm; id=1.57 mm; od=3.18 mm) containing either 100 μg/ml E2 (Sigma Chemical Co., St Louis, MO) in sunflower oil (OVX+E2 group) or oil vehicle (OVX+Veh group) (2). Four days later, the mice were anesthetized and sacrificed between 0900-1100h. Recent ELISA studies in our laboratory with the ES180S-100 CalBiotech Kit (43) established that this E2 treatment produces 7.59±0.7 pg/ml (Mean±SEM) serum E2 concentration in adult OVX female mice which falls within the high physiological range of intact cycling females (∼6pg/ml in diestrus and ∼8pg/ml in proestrus) (48). This 4-day E2 treatment also caused a robust 7-8-fold increase in uterine weight of OVX+E2 mice *vs.* the OVX+Veh controls (43).

### Gonadectomy of juvenile mice

To address the need of sex steroid exposure for the developmental induction of KP_LS_ neurons, juvenile mice between PND 21-31 were anesthetized with ketamine (25 mg/kg)/xylazine (5 mg/kg)/promethazine (2.5 mg/kg) and gonadectomized using sterile surgical techniques. 30 µg/10 g bw Augmentin dissolved in saline was injected in the peritoneal cavity of females. Wounds both in females and males were clipped and treated with Baneocin ointment to prevent infections. At PND 70-80, all animals were sacrificed by transcardiac perfusion with 4% paraformaldehyde and the brains were processed for microscopic analysis of KP_LS_ neurons as described in other experiments.

### Steroid induction of KP_LS_ neurons in juvenile mice

To induce new KP_LS_ neurons with a 4-day E2 treatment in juvenile (PND 20) females (N=4) and males (N=3), the same silastic capsules with E2 were used as the ones described above for treatment of adult mice. Other juvenile male animals were implanted for 4 days with another type of silastic capsule (Sanitech, Havant, UK; l=10 mm; id=1.47 mm; od=1.96 mm) that was filled with T (N=4) or DHT (N=4) powder in an attempt to advance the occurrence of KP_LS_ neurons. The mice were sacrificed by transcardiac perfusion with 4% paraformaldehyde after 4 days of steroid treatment. Tissue processing and cell counting with fluorescent microscopy were carried out as described in developmental studies.

### Detection of estrogen receptor-**α** in KP_LS_ neurons

Twenty-µm-thick paraformaldehyde-fixed histological sections of female mice (N=3) were used to study the presence of ERα immunoreactivity in the cell nuclei of KP_LS_ neurons recognized by their green fluorescence. For pretreatment, the sections were thoroughly rinsed in PBS and incubated with a mixture of 1% H_2_O_2_ and 0.5% Triton X-100 for 20 min. To maximize the immunofluorescent labeling, tyramide signal amplification (TSA) was carried out using the following incubation steps: monoclonal ERα antibodies (MA5–13191; Thermo Fisher Scientific; 1:500; 12h; RT), biotin-conjugated anti-mouse IgG (Jackson ImmunoResearch Laboratories; 1:500; 2h; RT), ABC Elite reagent (Vector; 1:1000 in 0.05M Tris-HCl buffer; 30 min; RT), Cy3-tyramide (diluted 1:1000 with 0.05M Tris-HCl buffer, pH 7.6, containing 0.003% H_2_O_2_; 30 min; RT). Negative control sections processed as above with the omission of the primary antibodies remained entirely devoid of the Cy3 labeling for ERα.

### Identification of cholinergic input to KP_LS_ neurons

Following the same tissue pretreatment and immunofluorescence strategies as described above, septum sections of diestrus female mice (N=3) containing KP-Cre/ZsGreen neurons were processed to detect the cholinergic marker vesicular acetylcholine transporter (VAChT) through the following steps: Guinea pig polyclonal VAChT antibody (#139 103; Synaptic Systems; 1:1000; 12h; RT), biotin-conjugated anti-guinea pig IgG (Jackson ImmunoResearch Laboratories; 1:500; 2h; RT), ABC Elite reagent (Vector; 1:1000 in 0.05M Tris-HCl buffer; 30 min; RT), Cy3-tyramide (diluted 1:1000 with 0.05M Tris-HCl buffer, pH 7.6, containing 0.003% H_2_O_2_; 30 min; RT). Sections processed with the omission of the primary antibodies remained devoid of Cy3 labeling.

### Fluorescent *in situ* hybridization

Fluorescent *in situ* hybridization detection of *Cartpt* and *Gad1* mRNAs with digoxigenin-labeled cRNA hybridization probes (targeting bases 61-466 of NM_013732 and bases 317–892 of NM_008077.4, respectively) was modified from previous procedures (49). To maintain ZsGreen fluorescence, slide-mounted 12-µm-thick sections of the septum were prepared from the brain of OVX+E2 (N=3) and OVX+Veh (N=3) mice perfused with 4% paraformaldehyde, instead of using the fresh-frozen tissue paradigm. Following posthybridization, fluorescent visualization of the nonisotopic signals used anti-digoxigenin antibodies conjugated to horseradish peroxidase (anti-digoxigenin-POD; Fab fragment; 1/200; Merck). This was followed by the deposition of biotin tyramide (1:1000; 30 min), and incubation of the sections in avidin-Cy3 (Jackson ImmunoResearch; 1:1000; 1h).

### Light and fluorescent microscopy

For fluorescent microscopy, sections were mounted from 0.1M Tris-HCl buffer (pH 7.6), air-dried and coverslipped with the aqueous mounting medium Mowiol. Images were captured with a Zeiss AxioImager M1 microscope (Carl Zeiss, Göttingen, Germany). Fluorescent images were analyzed using filter sets for fluorescein isothiocyanate or rhodamine. Digital images were captured with an AxioCam MRc5 digital camera (Carl Zeiss) and the AxioVision Se64 Rel.4.9.1 software.

#### Confocal microscopy

Cholinergic inputs to KP_LS_ neurons were studied with a Zeiss LSM780 confocal microscope using the Zen software (CarlZeiss). ZsGreen was detected with the 488 nm and Cy3 with the 561 nm laser line. Emission filters were 493–556 nm for ZsGreen and 570–624 nm for Cy3. Emission crosstalk between the fluorophores was prevented using ‘smart setup’ function. The final figures were adjusted in the ZEN Black Edition software (Carl Zeiss, Göttingen, Germany) using the magenta-green color combination and saved as TIF files.

### Slice electrophysiology

#### Brain slice preparation

Brain slices were prepared as described earlier (50) with slight modification. Briefly, OVX+Veh or OVX+E2 adult KP/ZsGreen female mice (36) were decapitated in deep isoflurane anesthesia. The brain was immersed in ice-cold low-Na cutting solution bubbled with carbogen (mixture of 95% O_2_ and 5% CO_2_). The cutting solution contained the following (in mM): saccharose 205, KCl 2.5, NaHCO_3_ 26, MgCl_2_ 5, NaH_2_PO_4_ 1.25, CaCl_2_ 1, glucose 10. Brain blocks including the lateral septum (LS) were dissected. 220-μm-thick coronal slices were prepared with a VT-1000S vibratome (Leica Microsystems) and transferred into oxygenated artificial cerebrospinal fluid (aCSF; 33°C) containing (in mM): NaCl 130, KCl 3.5, NaHCO_3_ 26, MgSO_4_ 1.2, NaH_2_PO_4_ 1.25, CaCl_2_ 2.5, glucose 10. The solution was then allowed to equilibrate to room temperature for 1 hour.

#### Whole cell patch clamp experiments

Recordings were carried out in oxygenated aCSF at 33°C using Axopatch-200B patch-clamp amplifier, Digidata-1550B data acquisition system, and pClamp 10.7 software (Molecular Devices Co., Silicon Valley, CA, USA). Neurons were visualized with a BX51WI IR-DIC microscope (Olympus Co., Tokyo, Japan). KP-ZsGreen neurons showing green fluorescence in the lateral septum (KP_LS_) were identified by a brief illumination (CoolLED, pE-100, Andover, UK) at 470 nm using an epifluorescent filter set. The patch electrodes (OD=1.5 mm, thin wall; WPI, Worcester, MA, USA) were pulled with a Flaming-Brown P-97 puller (Sutter Instrument Co., Novato, CA, USA). Electrode resistance was 2–3 MΩ. The intracellular pipette solution contained (in mM): K-gluconate 130, KCl 10, NaCl 10, HEPES 10, MgCl_2_ 0.1, EGTA 1, Mg-ATP 4, Na-GTP 0.3. pH=7.2-7.3 with KOH. Osmolarity was adjusted to 300 mOsm with D-sorbitol. Pipette offset potential, series resistance (R_s_) and capacitance were compensated before recording. Only cells with low holding current (< ≈50 pA) and stable baseline were used. Input resistance (R_in_), R_s_, and membrane capacitance (C_m_) were also measured before each recording by using 5 mV hyperpolarizing pulses. To ensure consistent recording qualities, only cells with R_s_<20 MΩ, R_in_>300MΩ, and C_m_ >10 pF were accepted.

Spontaneous firing activity and resting potential of KP_LS_ neurons were recorded in whole-cell current clamp mode at 0 pA holding current. Measurements started with a control recording (1 min). Then a single bolus of muscarinic ACh-R agonist muscarine (100 μM) (51) was pipetted into the aCSF-filled measurement chamber, respectively, and the recording continued for a further 7 min.

Pre-treatment with the extracellularly applied selective antagonist of the M2 subtype of mACh-R methoctramine (2 μM, Sigma) (52) or the voltage-gated Na-channel blocker tetrodotoxin (TTX, 660 nM, Tocris) started 10 min before adding muscarine. These inhibitors were continuously present in the aCSF during the electrophysiological recording.

Each neuron served as its own control when drug effects were evaluated.

#### Statistical analysis

Recordings were stored and analyzed off-line. Event detection was performed using the Clampfit module of the pClamp 10.7 software (Molecular Devices Co., Silicon Valley, CA, USA).

Mean firing rates were calculated as the number of action potentials (APs) divided by the control and treatment time periods, respectively. All the recordings were self-controlled in each neuron and the effects were expressed as percentage changes relative to the control periods. Duration of the effect of muscarine was defined as the period from the point all the APs disappeared till appearance of the first AP after the silent phase.

Treatment group data were expressed as mean ± SEM. Statistical significance was determined with two-tailed paired and unpaired Student’s *t*-tests and One-way ANOVA followed by Dunnett’s post-test using the Prism 3.0 software (GraphPad Inc., CA). Differences were considered significant at *p<*0.05.

### Immunohistochemical detection of kisspeptin in the human septum

#### Human tissues

Adult human brain tissues (N=3) from 38, 39 and 53 year-old male individuals without known neurological disorders were collected from autopsies at the 1st Department of Pathology and Experimental Cancer Research, Semmelweis University, Budapest, Hungary. Ethic permissions were obtained from the Regional and Institutional Committee of Science and Research Ethics of Semmelweis University (SE-TUKEB 251/2016), in accordance with the Hungarian Law (1997 CLIV and 18/1998/XII.27. EÜM Decree/) and the World Medical Association Declaration of Helsinki. The septum was dissected, cut into 4 mm-thick slices in the coronal plane, immersion-fixed with buffered 4% paraformaldehyde for 48 h, infiltrated with 20 % sucrose, embedded in Jung tissue freezing medium (Leica Biosystems, Nussloch, Germany) and snap-frozen on powdered dry ice. Then, 30-μm-thick coronal sections were collected with a Leica SM 2000R freezing microtome into tissue culture plates filled with anti-freeze solution (30% ethylene glycol; 25% glycerol; 0.05 M phosphate buffer; pH 7.4) and stored at -20 °C.

#### Immunohistochemistry

The sections were rinsed in PBS, pretreated with a mixture of 1% H_2_O_2_ and 0.5% Triton X-100 for 20 min and subjected to antigen retrieval with 0.1 M citrate buffer (pH 6.0) at 80 °C for 30 min. Every 24th section was immunostained with a well-characterized sheep polyclonal antiserum (GQ2; 1:75,000) against human kisspeptin-54 (aa 68-121 of NP_002247.3) (9, 53) using biotinylated secondary antibodies (Jackson ImmunoResearch Europe, Cambridgeshire, UK; 1:500), the ABC Elite reagent (Vector, Burlingame, CA; 1:1000; 1h) and the nickel-intensified diaminobenzidine chromogen [10 mg diaminobenzidine, 30 mg nickel-ammonium-sulfate and 0.003% H_2_O_2_ in 24 ml Tris-HCl buffer solution (0.05M; pH 8.0)]. Immunostained sections were mounted on 75 mm X 50 mm microscope slides from 0.3% polyvinyl alcohol, air-dried, dehydrated with 70%, 95% and 100% ethanol (5 min each), cleared with xylenes (2X5 min) and coverslipped with DPX mounting medium (Merck, Darmstadt, Germany).

### Transcriptomic studies

#### Section preparation for LCM

For all experiments, reagents were of molecular biology grade. Buffers were pretreated overnight with diethylpyrocarbonate (DEPC; Merck; 1 ml/L) and autoclaved or prepared using DEPC-treated and autoclaved water as diluent. The working area was cleaned with RNaseZAP (Merck KGaA, Darmstadt, Germany). To minimize possible adverse effects of fixation on RNA integrity and recovery, 0.5% formaldehyde in 0.1 M PBS (80 ml) was used for transcardiac perfusion of OVX+Veh (N=9) and OVX+E2 (N=12) KP-Cre/ZsGreen mice, followed by ice-cold 20% sucrose in PBS (50 ml). The brains were snap-frozen on pulverized dry ice and stored at -80 °C until sectioned with a cryostat. Twelve-µm-thick sections containing the LS were thaw-mounted onto PEN membrane glass slides (Membrane Slide 1.0 PEN, Carl Zeiss), air-dried in the cryostat chamber and preprocessed for LCM as reported elsewhere (25, 43). In brief, the slides were immersed sequentially in 50% EtOH (20 sec), n-buthanol:EtOH (25:1; 90 sec) and xylene substitute:n-butanol (25:1; 60 sec). Then, they were stored at -80 °C in slide mailers with silica gel desiccants or processed immediately for LCM.

#### Laser capture microdissection

Fluorescently tagged KP neurons (N=360/library) were microdissected individually and pressure-catapulted into 0.5 ml tube caps (Adhesive Cap 200, Carl Zeiss) with a single laser pulse using a 40× objective lens and the PALM Microbeam system and RoboSoftware (Carl Zeiss). Each neuronal pool was collected proportionally from 3 brains and stored in LCM tube caps at -80 °C until RNA extraction.

#### RNA extraction

RNA samples were prepared with the Arcturus Paradise PLUS FFPE RNA Isolation Kit (Applied Biosystems, Waltham, MA, USA).

#### RNA-Seq library preparation

Sequencing libraries were prepared with the TruSeq Stranded Total RNA Library Preparation Gold kit (Illumina, San Diego, CA, USA), using 16 PCR cycles for cDNA fragment enrichment. cDNA samples were subjected to Bioanalyzer analysis and a 1 nM library mix (20 µl) containing proportionally the indexed libraries was sequenced with an Illumina NextSeq500 instrument using the NextSeq500/550 High Output v2.5 kit (75 cycles).

#### Bioinformatics

Trimmomatic 0.39 (settings: LEADING:3, TRAILING:3, SLIDINGWINDOW:4:15, MINLEN:36) (54) and Cutadapt 4.4 (55) were used to remove low quality and adapter sequences, respectively. Remaining reads were mapped to the Ensembl mm109 mouse reference genome using STAR (v 2.7.10b) (56). Read assignment to genes, read summarization and gene level quantification were performed by featureCounts (subread v 2.0.6) (57). The counts per million read (CPM) values were calculated with the edgeR R-package (58).

#### Assessing KP cell type similarities

Transcripts with the highest mean CPMs were identified in KP_LS_, KP_ARC_ and KP_RP3V_ transcriptomes using functional categories of the KEGG BRITE database of the Kyoto Encyclopedia of Genes and Genomes; KEGG) (59) and a Neuropeptide database (http://www.neuropeptides.nl/).

#### Differential expression analysis

Differential expression analysis was performed with the DESeq2 R-package (60), following correction for unwanted variations with RUVSeq (61). Differences in mRNA expression levels were quantified by log_2_ fold changes (log_2_FC). To take multiple testing into account, p values were corrected by the method of Benjamini and Hochberg (62). Volcano plot and heat map, made with the EnhancedVolcano and Pheatmap program packages, respectively, were used to illustrate all transcripts that were expressed differentially.

## Acknowledgements

This work was supported by grants to Project no. RRF-2.3.1-21-2022-00011, titled National Laboratory of Translational Neuroscience implemented with the support provided by the Recovery and Resilience Facility of the European Union within the framework of Programme Széchenyi Plan Plus, the National Research, Development and Innovation Office (K128317 and 138137 to E.H. and PD134837 to K.S) and the Hungarian Research Network (SA-104/2021) of Hungary.

## Data availability

RNA sequencing files for KP_LS_ neurons are available in BioProject with the accession number PRJNA1017786. Re-analyzed sequencing files for KP_ARC_ and KP_RP3V_ neurons are cited in original publications (25, 43).

## Code availability

Scripts will be available upon request.

## Supporting information

This article contains **Table S1-4** as supporting information.

## CRediT author statement

**Conceptualization**, S.S., B.G., S.T., M.S., I.F., K.S., É.R., S.P., G.W., C.F., E.H.

**Methodology,** S.S., B.Go., S.T., M.S., I.F., K.S., S.P., G.W., E.H.

**Investigation,** S.S., B.G., S.T., M.S., I.F., K.S., S.P., G.W., E.H.

**Writing – editing,** I.F., M.S., E.H.

**Methodology,** S.S., B.G., S.T., M.S., I.F., K.S., S.P., G.W., E.H.

**Funding acquisition and supervision,** K.S., E.H.

## Ethics declaration

Animal experiments were carried out in accordance with the Institutional Ethical Codex, Hungarian Act of Animal Care and Experimentation (1998, XXVIII, section 243/1998) and the European Union guidelines (directive 2010/63/EU) and approved by the Animal Care and Use Committee of the Institute of Experimental Medicine. All efforts were made to minimize potential pain or suffering and to reduce the number of animals used.

For immunohistochemical studies of human *postmortem* brains, ethic permissions were obtained from the Regional and Institutional Committee of Science and Research Ethics of Semmelweis University (SE-TUKEB 251/2016), in accordance with the Hungarian Law (1997 CLIV and 18/1998/XII.27. EÜM Decree/) and the World Medical Association Declaration of Helsinki.

## Competing Interests

The authors declare no competing interests.

## Notes

### Competing Interest Statement

The authors have declared no competing interest.

